# Timescales of adaptation to context in horizontal sound localization

**DOI:** 10.1101/2022.11.27.518080

**Authors:** Gabriela Andrejková, Virginia Best, Norbert Kopčo

**Affiliations:** Institute of Computer Science, Faculty of Science, P. J. Šafárik University, Jesenná 5, Košice, 04001, Slovakia; Department of Speech, Language, and Hearing Sciences, Boston University, Boston, MA 02215, USA

## Abstract

Psychophysical experiments explored how the repeated presentation of a context, consisting of an adaptor and a target, induces plasticity in the localization of an identical target presented alone on interleaved trials. The plasticity, and its time course, was examined both in a classroom and in an anechoic chamber. Adaptors and targets were 2-ms noise clicks and listeners were tasked with localizing the targets while ignoring the adaptors (when present). The context had either a fixed temporal structure, consisting of a single-click adaptor and a target, or its structure varied from trial to trial, either containing a single-click or an 8-click adaptor. The adaptor was presented either from a frontal or a lateral location, fixed within a run. The presence of context caused responses to the isolated targets to be displaced up to 14° away from the adaptor location. This effect was stronger and slower if the context was variable, growing over the 5-minute duration of the runs. Additionally, the fixed-context buildup had a slower onset in the classroom. Overall, the results illustrate that sound localization is subject to slow adaptive processes that depend on the spatial and temporal structure of the context and on the level of reverberation in the environment.

## I. INTRODUCTION

Auditory spatial perception is highly adaptive (Carlile, 2014; King et al., 2000). Changes in horizontal sound localization can be induced by visual stimulation (Recanzone, 1998), feedback training (Klingel et al., 2021; Shinn-Cunningham et al., 1998), a change in the acoustic environment (Shinn-Cunningham et al., 2005), alterations in the mapping between acoustic cues and source locations (Kumpik et al., 2010; Trapeau & Schoenwiesner, 2018; van Wanrooij & van Opstal, 2007), or by other stimuli presented either simultaneously with the target (Braasch & Hartung, 2002), or preceding the target (Kopčo et al., 2010). The adaptation induced by preceding stimulation has been observed on long time scales of tens of seconds and minutes, e.g., in the auditory localization aftereffect induced by prolonged presentation of an adaptor (Carlile et al., 2001; Phillips & Hall, 2005; Thurlow & Jack, 1973), or in the precedence buildup induced by repeated presentation of ‘lead-lag’ stimulus pairs (Djelani & Blauert, 2001; Freyman et al., 1991). Studies of auditory localization aftereffects typically used a long continuous adaptor immediately followed by a target (Carlile et al., 2001; Thurlow & Jack, 1973), or even overlapping with the target (Canévet & Meunier, 1996). They observed a repulsion by the adaptor, i.e., biases in the perceived target locations away from the adaptor location. Here, we examine an adaptive effect qualitatively similar to the localization aftereffect but induced by the trial-to-trial acoustic “context” in which target sounds are presented. In our experiments, the target is a 2-ms broadband noise burst (referred to here as a ‘click’) (Kopčo et al., 2007). On some trials it is immediately preceded by an identical adaptor click (or clicks), and on other trials it is presented in isolation. Of interest here are localization biases for the target-only trials that are induced when those trials are randomly interleaved with adaptor-target trials. This effect, called *contextual plasticity* (CP), was observed in our previous work as repulsive biases of up to 10° in localization of the single-click targets.

Several different mechanisms have been proposed as underlying localization biases. First, some adaptation or fatiguing of the peripheral neural representation due to prolonged stimulation is often assumed (Carlile et al., 2001; Flugel, 1921). Second, a rebalancing of the putative hemispheric channels subserving spatial processing in humans has been proposed (Dingle et al., 2012; Phillips & Hall, 2005). Third, recent models based on known physiology of subcortical binaural circuits suggest that adaptation in response to the preceding context causes a rescaling of the spatial representation with the goal of increasing perceptual spatial separability of frequently presented sounds at the cost of inducing localization biases (Dahmen et al., 2010; Lingner et al., 2018; Maddox et al., 2014). Finally, an active centrally driven suppression of reverberation has been proposed for the precedence buildup, a potentially related phenomenon (Clifton et al., 2002).

The current study is the fourth in a series that examines CP. The original study (Kopčo et al., 2007) reported CP as an unexpected effect observed both in anechoic and reverberant rooms. Kopčo et al., (2015) showed that the effect is driven by adaptation in auditory perceptual representations as opposed to motor response-related representations, as it was observed for various response methods and with or without visual inputs. Finally, Hládek et al., (2017) showed that the strength of CP depends on the number of adaptor clicks and their similarity to the target. The goal of this fourth study is to examine how variability in the context affects CP and to present a detailed analysis of the temporal profile of CP.

Our analysis is based on data from two experiments using an identical design: one performed in a small classroom (Exp. 1) and one performed in an anechoic chamber (Exp. 2). While the experiments were primarily designed to examine the fast adaptation effects of the immediately preceding adaptors on timescales shorter than 0.5 secs (these data were reported in (Kopčo et al., 2007, 2017), the current study only focuses on the slower effects related to CP (some of which were reported in the previous studies without detailed analysis). In the experiments, CP was induced by context trials in which the adaptor was located either in front of or to the side of the listener (Fig. 1A), in one of two stimulus conditions (Fig. 1B): in the *fixed context* condition, the adaptor always contained one click (Kopčo et al., 2007), while in the *variable context* condition, the adaptor was either a single click or a train of 8 clicks, varying from trial to trial (Kopčo et al., 2017).

**FIG. 1.**
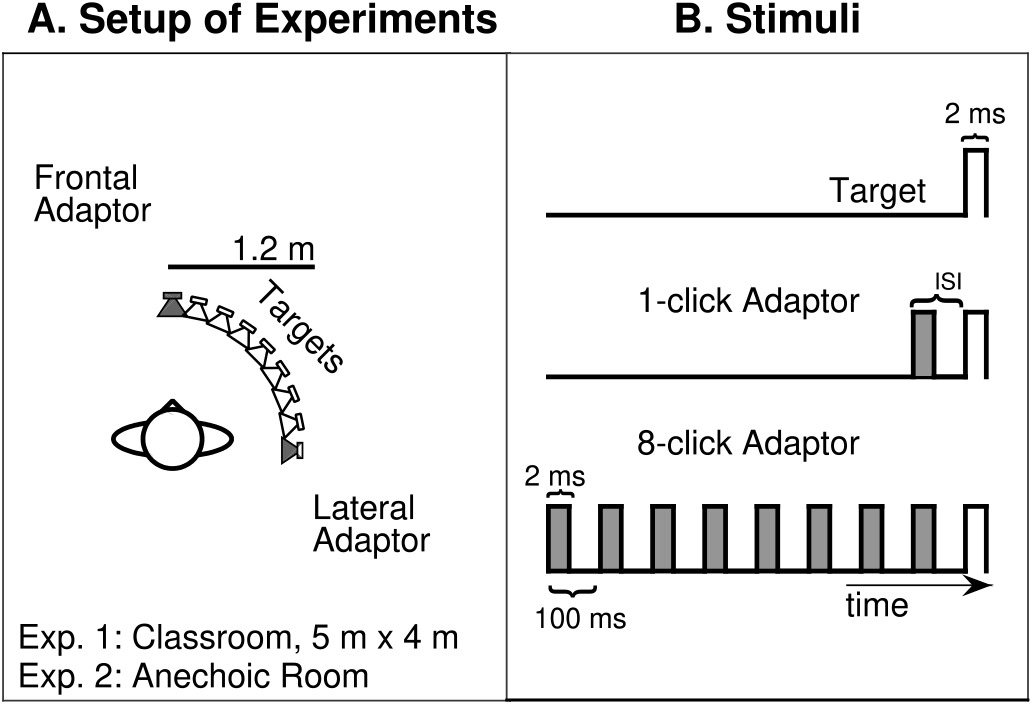
Experimental setup and stimuli. A. Arrangement of the loudspeaker array (shown here on the subject’s right-hand side). The adaptor (loudspeaker in grey color) was in the frontal position for half of the runs and in in the lateral position for the other half. B. Temporal structure of the target-only and adaptor-target stimuli, with adaptor in grey and target in white. Inter-stimulus interval, the time interval between the final adaptor click onset to the target click onset, ranged from 25 to 400 ms.

We addressed several questions related to the time course of CP. First, while we expected that CP would be stronger in the variable context condition as the average number of context clicks is higher in this condition (Hládek et al., 2017), we tested the hypothesis that it might also be slower to asymptote as the context varies from trial to trial. We also hypothesized that CP may be weaker and/or slower in the reverberant environment, as reverberation tends to make the spatio-temporal distribution of stimuli more uniform, which may reduce the strength of adaptation. Finally, we examined whether CP has a fast component on the time scale of seconds, observable when the context is varying from trial to trial.

## II. METHODS

The data described here were collected as part of two experiments previously reported in (Kopčo et al., 2007, 2017). The subjects, environments, and stimuli are the same as in those studies, but are briefly described again here.

### A. Subjects

Seven listeners (three females) with ages ranging from 23 to 32 years participated in Exp. 1 (Classroom), and four of these listeners also participated in Exp. 2 (Anechoic Room). All listeners reported normal hearing and gave informed consent as approved by the Institutional Review Board.

### B. Setup and listening environment

Exp. 1 was conducted in an empty, quiet rectangular reverberant room. The reverberation times in octave bands centered at 500, 1000, 2000, and 4000 Hz were 613, 508, 512, and 478 ms, respectively. The background noise level was 39 dBA. Exp. 2 was conducted in an anechoic room. Nine loudspeakers (Bose Acoustimass, Bose, Framingham, MA) were positioned on an arc with diameter of 1.2 m spanning 90°. The listener was seated approximately in the center of either room with his/her head held stable by a headrest. He/she sat in the center of the arc and faced either the left-most loudspeaker (so that the targets occurred on his/her right, see Fig. 1A) or the right-most loudspeaker (setup mirror-flipped compared to Fig. 1A). In the following, 0° azimuth always represents the location directly ahead of the listener, and 90° is the location of the left-or right-most speaker (depending on the listener orientation). Digital stimuli were generated by a TDT System 3 audio interface and passed through power amplifiers (Crown D-75A, Crown Audio, Elkhart, IN) to the loudspeakers. The listener kept their eyes closed during experimental runs and held a pointer in one hand for indicating the perceived direction of each target. A Polhemus FastTrak electromagnetic tracker was used to measure the location of the listener’s head, the approximate location of the loudspeakers, and the listener’s responses.

### C. Stimuli and procedure

The target was a single 2-ms frozen noise burst (click) presented at 67 dBA (Fig. 1B). An identical click was used for the adaptor in the 1-click context trials. Eight such clicks presented at the rate of 10/sec (T = 100 ms) made up the adaptor in the 8-click context trials. Within a run, the context was either fixed or variable. In the fixed context runs, only the 1-click contexts were used, the ratio of contextual to target-only trials was 5:1, and the adaptor-target inter-stimulus interval, measured from the peak of the final adaptor click to the peak of the target click, was 25, 50, 100, 200, or 400 ms. In the variable context runs, the ratio of 8-click context to 1-click context to target-only trials was 2:2:1 and the inter-stimulus interval was 50 or 200 ms. On each trial, the target location was randomly selected from one of the seven central loudspeakers (spanning approximately 11°–79° azimuth), while the adaptor, if any, was played from a loudspeaker that was fixed within a run. Every combination of the six (fixed context) or five (variable context) trial types and seven target locations was presented four times in random order within a run, resulting in 168 trials in the fixed context runs and 140 trials in the variable context runs. The subject changed his/her orientation after each run to face either the left-most loudspeaker or the right-most loudspeaker by rotating his/her whole body.

Exps. 1 and 2 each comprised eight sessions, 4 for the fixed context and 4 for the variable context. Each session, which took approximately 30 min, contained four randomly ordered runs, one for each combination of subject orientation (facing the left-most speaker, facing the right-most speaker) and context adaptor location (frontal, lateral). The total duration of a run was relatively consistent, with across-subject means and standard deviations of 5.3±0.6 min (Exp. 1, fixed context), 5.1±0.7 min (Exp. 2, fixed context), 5.6±0.6 min (Exp. 1, variable context), and 5.3±0.5 min (Exp. 2, variable context).

### D. Data analysis

The current analyses focus exclusively on data from the target-only trials (see Kopčo et al., 2007, 2017) for the analysis of the context trial data). There were only small differences between the data sets collected with the two subject orientations, and thus the data were collapsed across the orientations and sessions and analyzed as if the subject always faced the leftmost loudspeaker. Since only a subset of the Exp. 1 subjects participated in Exp. 2, data are also presented for this subset of 4 subjects in Exp. 1, to allow a direct comparison of the effect of room across the subjects. To analyze the temporal profile of CP, data from each run were divided into 4 subruns, as each run contained 4 repetitions of each stimulus combination, presented in a pseudo-random order such that any combination was repeated only after all other combinations were presented. All reported statistical analyses were performed as multi-way repeated measures analyses of variance (ANOVAs), using CLEAVE software (Herron, 2005). The reported statistical values were corrected for potential violations of sphericity using the Greenhouse-Geisser epsilon.

## III. RESULTS

Three analyses are presented in the following sections. The first analysis focuses on the spatial profile of CP and its change over time (Section III.A). Then, the temporal profile of the CP is analyzed on time scales of minutes (Section III.B) and seconds (Section III.C).

### A. Spatial and temporal profiles of contextual plasticity

Fig. 2 shows the across-subject mean bias in localization responses as a function of target location, separately for the two context adaptor locations (circles for frontal vs. triangles for lateral), the two context conditions (red for fixed, blue for variable), and the two experiments (panels A and B for Exp. 1, C for Exp. 2). Panel B shows the Exp. 1 data for the 4 subjects who also participated in Exp. 2.

**FIG. 2.**
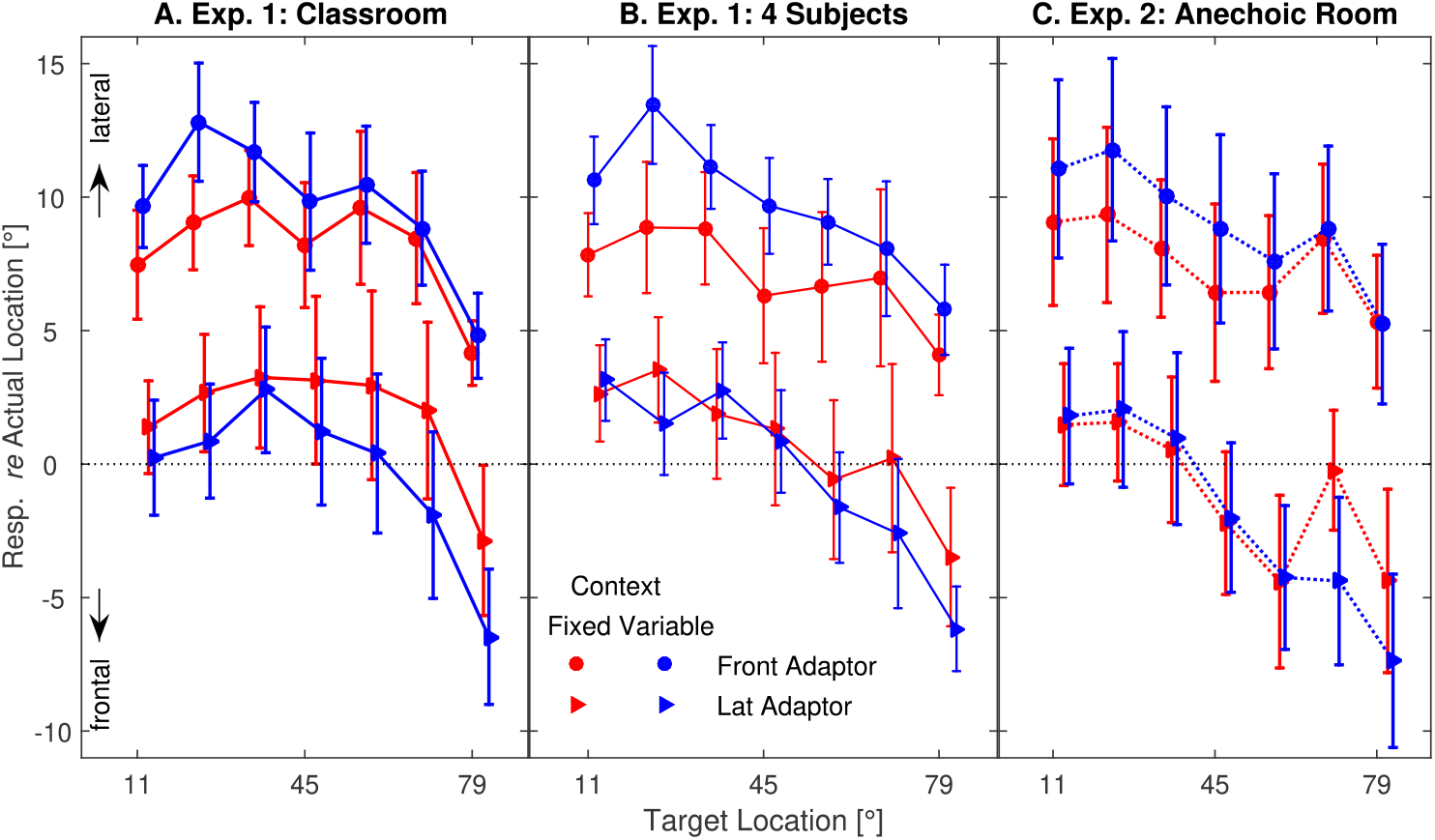
Mean response biases (+-SEM) in target-only trials in Exp. 1 (panel A) and Exp. 2 (panel C), plotted as a function of target location separately for each combination of context type and adaptor location. Panel B shows the Exp. 1 data for the 4 subjects who participated in Exp. 2.

To analyze the mean bias results, two ANOVAs were performed. The first ANOVA considered the Exp. 1 data on all 7 subjects (panel A), with factors of *Adaptor location* (frontal, lateral), *Target location* (7 locations from 11° to 79°), *Context type* (fixed, variable), and *Subrun* (1 to 4). This ANOVA found significant main effects of *Adaptor location* (F(1,6)=56.93, p=0.003, η _p_^2^=0.905) and *Target location* (F96,36)=4.76, p=0.0012, η _p_^2^=0.442), as well as significant interactions of *Target location x Subrun* (F(18,108)=2.24, p=0.0057, η _p_^2^=0.272), *Adaptor location x Subrun* (F(3,18)=27.52, p=0.0000, η _p_^2^= 0.821), *Context type x Adaptor location* (F(1,6)=11.49, p=0.0147, η _p_^2^=0.657), and *Context Type x Target location* (F(6,36)=3.95, p=0.0039, η _p_^2^=0.397). The second ANOVA considered both rooms and was restricted to the 4 subjects who performed the experiments in both rooms (panels B and C). It had an additional factor of *Room* (anechoic, reverberant), and it found significant interactions of *Context Type x Room x Subrun x Adaptor location* (F(3,9)=4.84, p=0.0285, η _p_^2^=0.617), *Context type x Target location* (F(6,18)=6.19, p=0.0012, η _p_^2^=0.674), *Context type x Adaptor location* (F(1,3)=12.54, p=0.0383, η _p_^2^=0.807), and *Subrun x Adaptor location* (F(3,9)=35.62, p=0.0000, η _p_^2^=0.922). No other main effects or interactions reached significance.

The data in Fig. 2 indicate that localization responses were biased relative to the actual target locations. The frontal context data (circles) were biased laterally by approximately 5 to 13°, while the lateral context data (triangles) were biased by −5 to 5°. Such “global” response biases are common in localization experiments and arise from a combination of factors including the response method (Kopčo et al., 2015). Of more interest here are differences in the bias depending on the context. The clearest effect shown in Fig. 2 is that the responses with frontal contexts are always biased more laterally than the responses with lateral contexts (triangles fall under circles in all three panels, confirmed by the main effect of *Adaptor location* in Exp. 1). This effect is overall stronger for the variable context than the fixed context, particularly near the adaptor locations (blue circles are above the red circles especially for the targets at 11-33°; blue triangles are below the red triangles especially for the targets at 56-79°; significant *Context type x Adaptor location* and *Context type x Target location* interactions). Because this pattern is approximately symmetric and complementary (dominated by the frontal adaptor for frontal targets and the lateral adaptor for lateral targets), the differences between frontal and lateral adaptor contexts are approximately target-location independent (corresponding red lines are approximately parallel, as are the corresponding blue lines; *Context type x Target location x Adaptor location* interaction is not significant).

Before comparing the results across the rooms, note that the results in panels A and B are very similar, i.e., that the subgroup of participants who also participated in Exp. 2 is representative of the larger group. Panels B and C show that the effect of context was also modulated by the room in which the stimuli were presented, and the ANOVA further suggests that the room effect changed over time (4-way *Context type x Room x Subrun x Adaptor location* and 2-way *Subrun x Adaptor location* interactions). These interactions did not include the *Target location* factor, suggesting again that the important features of CP are approximately target-location independent.

We operationalize CP in terms of CP_diff_ which is the difference between frontal and lateral context biases, averaged across target location. CP_diff_ is plotted as a function of subrun in Fig. 3A for both experiments (differentiated by line styles, which match Fig. 2) and contexts (line color) (Footnote 1). The results show that, overall, CP had a fast onset, reaching values between 3° and 7° within the first subrun. It continued to grow on the time scale of minutes in both conditions (top scale in Fig. 3A), with the rate of growth dependent on the context type and on the room. Overall, CP tended to be larger with the variable context (blue lines are above red lines) and in the anechoic room (dashed lines tend to be above solid lines). However, these effects varied over time and did not combine additively. Specifically, for the fixed context (red lines) the room effect (dashed vs. solid line) was the largest at the beginning of the run, while for the variable context (blue lines) it was largest at the end of the run. Finally, the variable context CP in both environments continued to grow between the 3^rd^ and 4^th^ subruns, suggesting that it did not reach its maximum over the 5-minute course of individual runs in this condition (reaching 12-14°). In the fixed context condition, CP appeared to reach its maximum of 8-10° by subrun 3.

**FIG. 3.**
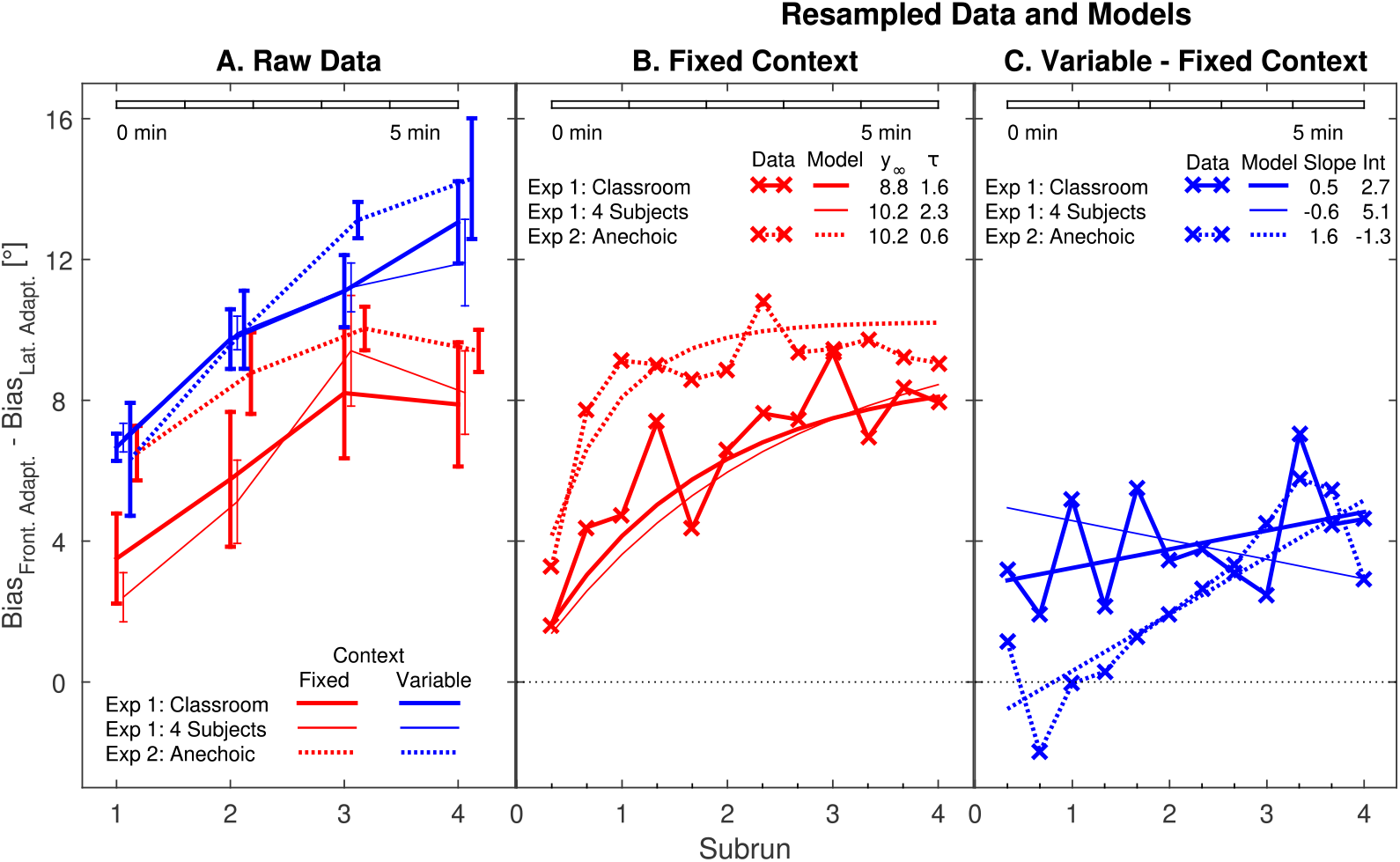
Temporal profile of CP_diff_ for the two contexts and two rooms. A. Mean CP_diff_ (±SEM) divided into 4 subruns. B. Fixed-context data rearranged to increase temporal resolution and modeled using exponential fits (fitted parameters are shown in the inset). C. Variable context re. fixed context data rearranged and modeled using linear fits (parameters shown in the inset).

## Discussion

The analysis of the spatial properties of CP showed that 1) CP is observed as a repulsion of responses away from the adaptor location that decreases with separation between target and adaptor, and that 2) the effect is stronger in the variable context condition where the overall adaptor click rate is higher. These results are consistent with previous studies (Hládek et al., 2017; Kopčo et al., 2015) but extends the finding to lateral as well as frontal adaptors. Additionally, we find that variable and fixed context effects are similar both in their strength and spatial extent for the frontal and lateral adaptor when expressed as a function of distance from the adaptor, suggesting that the spatial representation in which CP is induced is approximately uniform, even though auditory spatial resolution decreases with azimuth (Hartmann & Rakerd, 1989).

The temporal analysis of CP showed that the effects of room and context type interact and are combined non-additively. Specifically, CP was strong already at the beginning of the run in both rooms for the variable context and in the anechoic room also for the fixed context, while being relatively weak in the classroom fixed context runs. Towards the end of the runs, CP became largely independent of the environment while differing strongly for the two contexts. Specifically, the variable context CP continued to grow even after approximately 5 minutes, while in the fixed context the CP reached its maximum after 2-3 minutes, consistent with previous studies which only used fixed context (Hládek et al., 2017). Thus, varying the context from trial-to-trial causes at least a doubling of the time it takes CP to reach its asymptote, resulting in CP that is stronger (12-14° by subrun 4) than that observed with fixed 1-click context (8-10°) or fixed 8-click context (9°, (Hládek et al., 2017)).

### B. Modeling of the temporal profile of contextual plasticity

To further increase the temporal resolution, we grouped the data from targets at 11°, 22° and 34° into one target “triplet” and data from targets at 56°, 67°, and 79° into another target triplet. By this rearrangement, the temporal resolution could be increased three-fold, as each of the original 4 subruns now contained 3 data points approximately evenly distributed across it. Then, we used exponential fits to analyze the buildup of CP in the fixed context runs, and linear fits to describe the additional buildup in the variable context runs. Specifically, each subject’s fixed context CP data were fitted parametrically using the first-order exponential equation

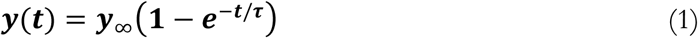

with time ***t*** in the units of subruns, yielding a time constant ***τ*** for the adaptation to the context (with **1/*τ*** as its rate) and a model estimate of the asymptotic value of CP, ***y***_∞_. The model assumed that the initial, pre-adaptation CP was 0 and that the asymptotic value of CP was equal for the two environments (consistent with the observed data). Thus only 3 parameters were fitted per subject, one ***y***_∞_ for both environments and one ***τ*** for each environment. The additional CP observed in the variable context (vs. fixed context) was modeled using a linear model as there was no evidence that the difference data deviated from linearity in either environment. The analysis was then focused on the estimated slope of the adaptation, which represents the temporal properties of the additional adaptation.

The results of this analysis are shown in Figs. 3B and 3C. In both figures, the mean data are shown by symbols ‘x’, the fits for the classroom are shown by thick solid lines (the 4-subject fit is shown by thin solid lines) and the fits for the anechoic room are shown by dotted lines.

The fixed context data and fits in Fig. 3B (red symbols and lines, corresponding to the red lines from panel A) show that the onset of CP is faster in the anechoic room than in the classroom, and that the difference between the anechoic and classroom data is only around 1° towards the end of the run. As mentioned above, given the small difference at the asymptote, the model was fitted such that only one common *y*_∞_ was used for both environments while τ values were separate. The common fitted value of *y*_∞_ was 10.2°. On the other hand, the time constant τ differed significantly between the environments for the 4 subjects who participated in both experiments. The mean τ was 2.3 subruns in the classroom and 0.6 subruns in the anechoic room (t(3)=-3.178, p=0.019).

Fig. 3C shows, for both environments, the difference between the variable and fixed context CP data (i.e., the difference between respective blue and red data from panel A), as well as the linear fits. The data show that the effect of variable context (re. fixed context) is approximately linear in both environments. In the classroom experiment, the variable context caused an additional repulsion from the adaptor location of approximately 4°, independent of time (solid lines). In the anechoic room, the effect of the variable context was much slower, growing from approximately 0° to 4°. A paired-samples t-test (t(3)= −4.7, p = 0.018) confirmed that the slopes of the fits were significantly different in the anechoic vs. reverberant room.

## Discussion

The modeling presented in section III.B confirmed the results of the behavioral data analysis of section III.A. The exponential model fitted to the fixed context data found a significant difference in adaptation rate between the two rooms, supporting the conclusion that the initial difference between the two rooms was mostly driven by a difference in speed, not strength, of CP, as the rate at least doubled in the anechoic room compared to the classroom.

The additional adaptation in the variable context showed either a constant or an approximately linear growth, uniform over the whole duration of the current runs. This again shows that the variable context, randomly switching between a 1-click and an 8-click adaptor, causes the adaptation to have a very slow component, much slower than those observed in our previous studies (e.g., (Hládek et al., 2017)) and resulting in a stronger CP. Note that the additional adaptation would likely have reached an asymptotic value if the runs were sufficiently long. However, since it did not reach its asymptote in the current experiment, and since the difference between the conditions was largely linear in both environments, a linear model was sufficient to describe the data.

Finally, note that the presented modeling always considered the difference between the frontal-adaptor and lateral-adaptor contexts, i.e., the CP^diff^, corresponding to a combination of two adaptive processes, one for each context. The Appendix provides the results of additional modeling performed separately for the two adaptor locations, which shows that the slow minute-scale adaptation correlates with the distribution of the stimuli in different contexts, consistent with the hypothesis that spatial auditory processing prioritizes discriminability of stimuli over unbiased localization (Lingner et al., 2018).

### C. Trial-to-trial adaptation in the variable context runs

The previous section showed that one effect of varying the context on a trial-to-trial basis was that the adaptation continued to evolve over the duration of an experimental run (around 5 minutes). Here, an analysis is performed on the time scale of individual trials, to examine 1) whether the extremely slow adaptation is accompanied and/or caused by a fast-varying plasticity changing after every context trial, and 2) whether this effect varies over the course of an experimental run. The variable context runs included two types of context trials (1-click and 8-click adaptors), randomly interleaved with the target-only trials. In this analysis, the target-only trials were split by the type of the preceding context trial (1-click or 8-click) and plotted as a function of subrun for the classroom (Fig. 4A) and the anechoic room (Fig. 4B). In the classroom, the trials preceded by an 8-click adaptor trial show a faster onset of CP, reaching the maximum of 12° by the second subrun (solid lines with asterisks), while the trials preceded by a 1-click adaptor trial show CP of around 6° in the first two subruns and only reach 12° adaptation in subrun 3 (lines with no symbols). On the other hand, in the anechoic room, CP grows throughout the run but there is no systematic difference based on the immediately preceding context trial type.

**FIG. 4.**
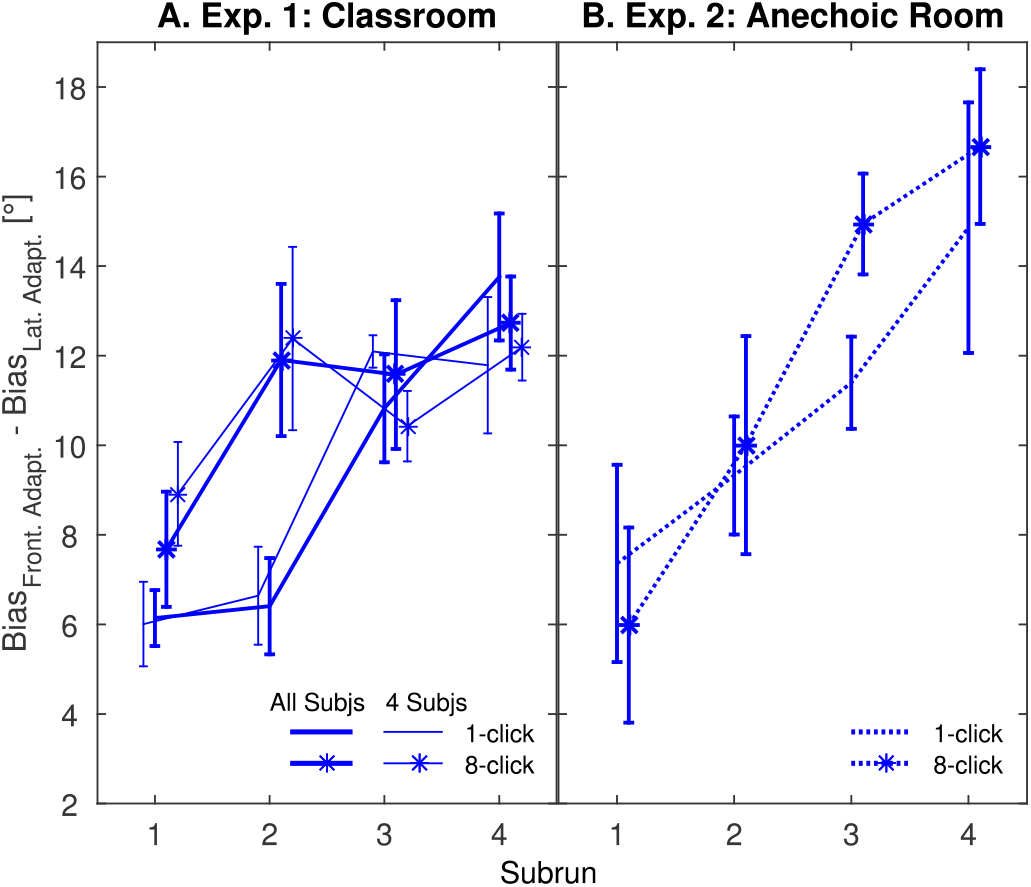
Effect of the context adaptor click rate (1-click vs. 8-click adaptor) in the immediately preceding trial on CP in the variable context condition. CP_diff_ is plotted as a function of subrun in the classroom (panel A) and the anechoic room (panel B).

Confirming these observations, an ANOVA with the factors of *Subrun* (1-4) and *Context* (1-click vs. 8-click) performed on the classroom data found a significant main effect of *Subrun* (F(3,18) = 14.93, p = 0.0003, η_p_^2^= 0.71) and a significant interaction *Context x Subrun* (F(3,18) = 3.59, p = 0.034, η _p_^2^= 0.37). A similar ANOVA performed on the anechoic data found no significant effects.

## Discussion

This analysis shows that, in some instances, CP was affected by the immediately preceding context trial, such that the effect was larger following 8-click vs. 1-click adaptors. This suggests a relatively fast contextual effect, corresponding to the 3-5 sec timescale of individual trials. However, it was only observed during the first half of each run and only in the classroom. One possible explanation of this effect is that, in addition to a slow adaptation, a reverberation-suppression mechanism related to precedence buildup (Brown et al., 2012) influences these trials. Specifically, it may be that reverberation suppression effects operating on the prolonged 8-click adaptor “spill over” to affect not only the target on that trial but also that on the subsequent trial. When only 1 adaptor click is presented in the context, this suppression is apparently reduced. It is not clear why this effect was restricted to the early part of each classroom run, though it is possible that the effect was simply not visible later as the CP saturated. An alternative mechanism might be related to perceptual organization, as the 8-click adaptor trials were designed to increase the perceptual segregation of the adaptors and targets (Kopčo et al., 2017). In any case, our data clearly show that each localization trial can be influenced by the immediately preceding trials, which may be an issue for task designs that intermix different conditions (e.g., Kopčo et al., 2017; Moore et al., 2020).

## IV. GENERAL DISCUSSION

Our work shows that the spatial and temporal distribution of stimuli (or “context”) in which a listener performs a localization task has a complex influence on their behavior. The main finding of the current study was that the repeated presentation of an adaptor-target context induced a slow adaptation in the localization of targets that 1) resulted in biases of up to 14° away from the adaptor location, 2) built up over at least 5 minutes, and 3) depended on the spatial and temporal structure of the adaptors, as well as on the presence of reverberation. Specifically, increasing the average number of adaptor clicks (variable context) resulted in a stronger CP, while both varying the number of clicks from trial to trial and an exposure to reverberation resulted in a slower temporal profile of the adaptation. Strikingly, the variable context resulted in adaptation that grew over time in both environments, resulting in the strongest CP we have observed to date, even stronger than that induced by a fixed 8-click context (9° observed in (Hládek et al., 2017)). These effects of context type and environment are likely due to some non-linear interaction of multiple adaptive processes that depend on the adaptor presentation rate, its variability, and the presence of reflections.

The spatial profile of CP was originally reported to be largely independent of the adaptor-target distance (Kopčo et al., 2007). Later studies, which only used a frontal adaptor and also included no-adaptor-baseline runs, showed that the effect is stronger near the adaptor location and that it largely disappears for targets separated by 80° from the adaptor (Hládek et al., 2017; Kopčo et al., 2015). The current study showed that the dependence of the CP strength on the separation from the adaptor also applies to the lateral adaptors, and that the repulsive effects of frontal and lateral adaptors are similar. It is worth noting, however, that the adaptor was always at the edge of the target range in the current study. It is possible that placing the adaptor in the middle of the target range and/or using targets symmetrically located around the midline, as in the previous localization aftereffect studies (Carlile et al., 2001; Phillips & Hall, 2001; Thurlow & Jack, 1973) would result in a different pattern of adaptation.

The main finding concerning the temporal profile was that varying the context from trial to trial produced CP that was very slow to stabilize, continuing to grow for at least 5 mins, while fixed context CP asymptoted after 1-2 mins (Hládek et al., 2017; Kopčo et al., 2015). Such extended adaptation has not been reported in previous localization aftereffect studies, which focused on effects occurring immediately post-adaptor (Carlile et al., 2001; Lingner et al., 2018), while other related studies likely observed such long-term adaptation but ascribed it to other factors (Moore et al., 2020). In future studies it would be very interesting to include long enough runs for CP to reach the asymptotic value in the variable context, so that it can be established how long such an adaptation can continue for. Another interesting finding was that, in the variable context, there was evidence for a fast adaptive component that is sensitive to the temporal structure of individual context trials. Since this fast component was not observed in the anechoic room, it is possible that it is related to reverberation suppression mechanisms evoked in the precedence effect and its buildup (Brown et al., 2012; Litovsky & Macmillan, 1994) or perceptual organization (Kopčo et al., 2017). On the other hand, our subject pool may have been too small to reveal similar effects in the anechoic room, and further investigations would be needed to make strong conclusions. Reverberation also affected the initial onset of CP, which was considerably slower in the classroom than in the anechoic room for the fixed context. Again, this difference may be related to precedence buildup mechanisms operating in the reverberant classroom. Overall, the effects of reverberation that we observed were small, and we did not find strong support for the hypothesis that CP would be weaker in reverberation where the presence of omnidirectional reverberation makes the distribution of energy more uniform around the listener.

Finally, while the data presented here are unable to distinguish between competing models of spatial adaptation that have been proposed in the literature, they provide some preliminary indications that may be worth following up on. For example, an exploratory analysis of the data (reported in the Appendix) shows that the slow component of CP can be well characterized as a linear drift in the spatial auditory representation in response to the overall spatial distribution of the stimuli in a particular run. Specifically, stronger drifts towards midline were observed with increased laterality of the distribution mean. Such a relationship is consistent with the idea that CP might be caused by adaptation of the neural representation that shifts it towards the stimulus distribution mean (Dahmen et al., 2010; Lingner et al., 2018). The specific neural mechanisms underlying the shift might include dynamic range adaptation (Dahmen et al., 2010), synaptic gain control (Stange et al., 2013), or re-balancing of excitatory and inhibitory inputs (Magnusson et al., 2008). This “shift” model offers an alternative to “suppression” models which posit that localization aftereffects are caused by local suppression/fatiguing of spatial channels near the adaptor (e.g., Carlile, 2014). Of course, it is possible that both shift and suppression mechanisms contribute to CP and related spatial adaptation phenomena. Future experiments specifically designed to untangle these mechanisms may bring further insights.

## ACKNOWLEGEMENTS

Work supported by the Slovak Scientific Grant Agency VEGA 1/0350/22 and by EU Danube Region Strategy grant ASH (Grant Nos. APVV DS-FR-19-0025, WTZ MULT 07/2020, 45268RE). VB was partially supported by NIH-NIDCD Award No. DC015760. The authors thank Bernhard Laback for his comments on an early version of this manuscript.

## APPENDIX

### Relationship between contextual plasticity buildup and stimulus distribution Motivation

In this exploratory analysis, we attempted to relate the temporal profile of CP to the spatial distribution of the stimuli in different contexts. Our goal here was to provide a preliminary test of competing hypotheses about the mechanisms underlying CP.

While the mechanism underlying CP is largely unknown, it shares many properties with the localization aftereffect (Phillips & Hall, 2005; Thurlow & Jack, 1973). Specifically, it results in similar shifts in the perceived target location away from the adaptor location, although on a longer time scale. Various models have previously been proposed for the localization aftereffect, many of them assuming that it is caused by some suppression in the neural representation of auditory space (Carlile, 2014; Dingle et al., 2012). It has also been suggested that the observed shifts are a result of a broad dynamic range adaptation of the auditory spatial representation, occurring when the stimulus distribution becomes concentrated in a subregion of the full horizontal spatial range (Dahmen et al., 2010). In this scenario, biases in responses are a negative side effect of the representation adapting to improve the spatial separability of targets presented within the subregion. This adaptation may be implemented by fitting the working point of the neural firing rates vs. the spatial location to the middle of the stimulus range (Lingner et al., 2018).

Motivated by the latter studies, here we examine the hypothesis that the auditory representation adapts to the non-uniform stimulus distribution in our experiments. We test a simplified prediction that the more skewed the stimulus distribution from the midline, the stronger the response biases induced by it. To test this prediction, we analyze the drifts in response biases over the course of individual runs from subrun 1 to subrun 4 and evaluate whether the slope of these drifts, averaged across target location, can be predicted by the size of the change in the stimulus distribution mean. The analysis focuses on the drifts, not on the absolute value of the change, because looking at the drifts 1) allows us to consider the frontal-context and lateral-context data separately, as we are only looking within a run, 2) only requires to use the 1^st^ subrun as a reference (no preadaptation reference was measured), and 3) allows the analysis to focus on the slow adaptation occurring on time scale larger than 1-2 minutes (i.e., the approximate duration of one subrun), in which the drifts were largely linear. Our analysis is performed on the data presented in the main body of the current paper, as well as on additional data from Kopčo et al., (2015).

#### 1. Data from current study

In Exps. 1 and 2, targets were presented from a frontal left or frontal right quadrant in the horizontal plane, with the adaptor always located at the edge of the target range (Fig. 1). The left-hand panel of Fig. A1 A shows the distributions of these click stimuli within a run, including both the adaptor and target clicks (bars), separately for the frontal-adaptor and lateral-adaptor runs (note that the distribution was identical in the two experiments). The symbols along the upper edge indicate the respective distribution means. In each of the four contexts, the stimuli are shown for the runs performed in the right-hand quadrant (the left-hand quadrant stimuli would add symmetrical distributions and means on the left-hand side). The distribution mean was between 9° for the frontal-adaptor variable-context runs (blue circle) and 81° for the lateral-adaptor variable context runs (blue triangle), with the respective fixed context means (red circle and triangle) falling between the variable context mean values. Based on our hypothesis, for the stimuli presented in the right-hand quadrant, the responses are expected to drift to the left, as the channels representing the left quadrant shift their receptive fields to the right. Additionally, this drift is expected to be larger in the lateral adaptor runs than in the frontal adaptor runs, as the distribution is skewed more positively (to the side) when the adapter is lateral.

The two left-hand panels of Fig. A1 B show, for the two experiments, the across-target average response bias as a function of subrun. The symbols represent the mean response bias in each subrun, separately for the fixed vs. variable contexts (red vs. blue), frontal vs. lateral adaptors (circles vs. triangles), and classroom vs. anechoic room (filled vs. open symbols). The lines show the across-subject average of linear fits through the data performed separately for each context (lines going through the triangle vs. circle data for the frontal vs. lateral context) and room (solid for classroom, dashed for anechoic room). The fits are very good, indicating that the adaptation is approximately linear over this time range. They have negative slopes for the lateral context (from −1.8 to − 0.94°/subrun) and slightly positive slopes for the frontal context (0.06 to 0.38°/subrun), an effect that tends to be stronger for the variable vs. fixed context (blue vs. red lines), especially in the anechoic room (dashed lines). These trends were confirmed by an ANOVA with factors of *Context type, Room, and Adaptor location*, performed on the slopes of the linear fits, which found a significant main effect of *Adaptor location* (F(1,3)=105.87, p=0.002, η _p_^2^= 0.9724) and a significant 3-way interaction (F(1,3)=10.43, p=0.048, η _p_^2^= 0.7766).

To directly evaluate the relationship between the distribution of the stimuli and the response drifts, the left-hand panel of Fig. A1 C plots the slope of the response drifts (from Fig. A1 B) as a function of the mean lateral position of the stimuli (from Fig. A1 A). There is a strong correlation, with the across-subject average *r* reaching 0.95 in the anechoic room and 0.86 in the classroom. A linear fit to the data (black line) shows that the slope of the drift in responses is inversely proportional to the mean of the stimulus distribution (slope of this fit is −0.033; t(6) = −10.4, p<0.0001). This general result is consistent with the idea that the drift occurs as a result of a dynamic range adjustment (Dahmen et al., 2010; Lingner et al., 2018). However, there is one aspect of the data that is not consistent with this idea. While the distribution means are all positive, predicting that the drift slopes should always be negative, the slopes of the drift for the frontal contexts are slightly positive (circles in Fig. A1 C). A potential explanation for this discrepancy is that, in addition to the distribution-dependent drifts, the responses also drifted due to some other factors, like a fatiguing of the motor responses, as the subjects used a hand-held pointer to respond.

#### 2. Data from Kopco et al. (2015)

To examine whether the slopes are influenced by the response method used by the listeners, we performed the same analysis on data from a previous study (Exp. 1 of (Kopčo et al., 2015)). That study was very similar to the current fixed-context classroom Exp. 1, differing only in two important aspects. First, three different response methods were used: 1) using a hand-held pointer while the eyes were closed (like in the current study), 2) using a hand-held pointer with the eyes open, and 3) a keyboard-based method that used vision but did not require any sensory-motor spatial transformation to respond. The right-hand panel of Fig. A1 A shows the stimulus distribution in this study. The frontal-adaptor runs had distributions very similar to Exp. 1 (green vs. red filled bars), while the baseline runs had a uniform distribution with a mean at 45° (black bars).

**FIG. A1.**
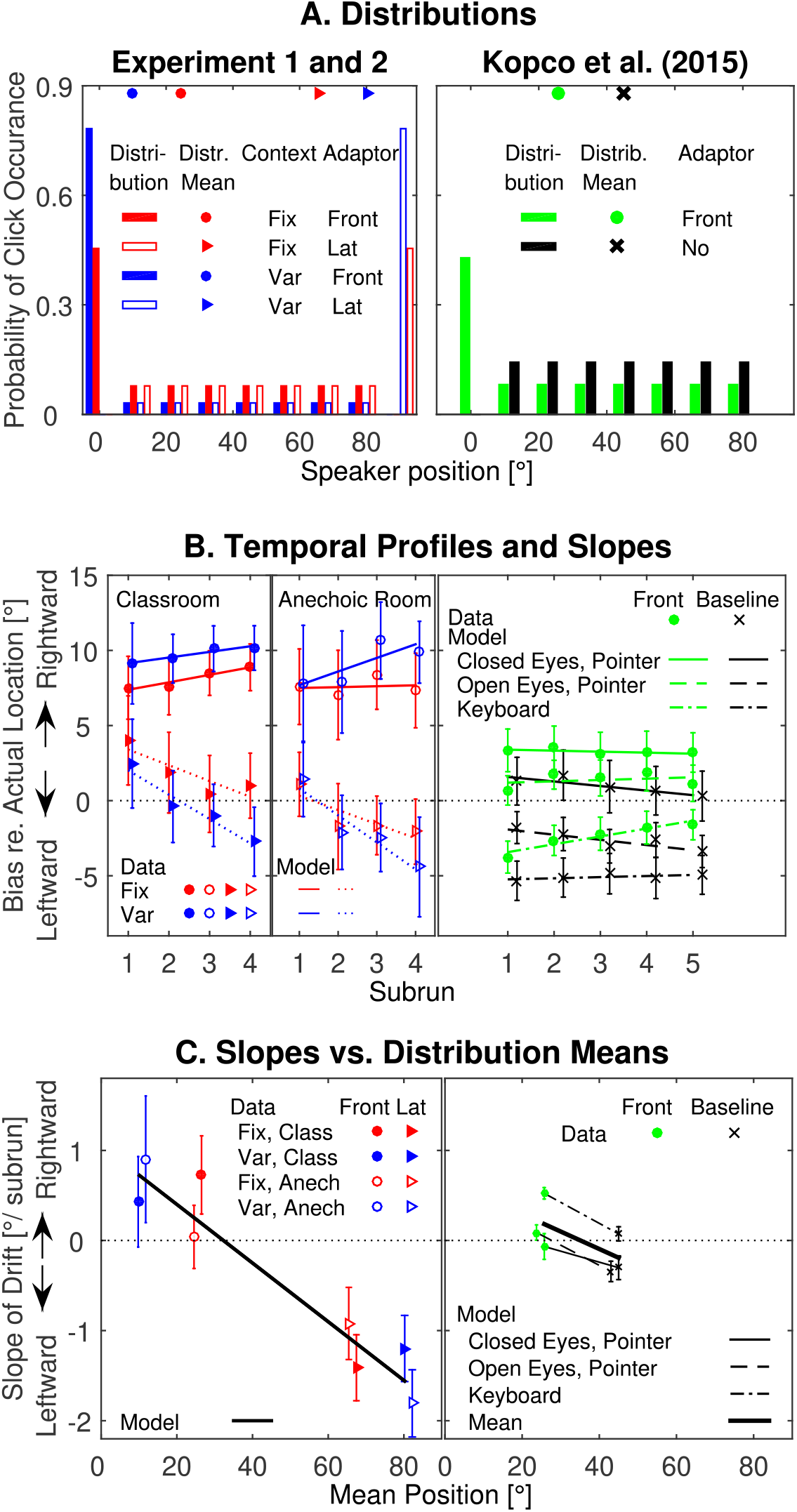
Relationship between stimulus distribution and drift in responses for current Experiments 1 and 2 (left-hand panels) and from a previous study ((Kopčo et al., 2015); right-hand panels). A. Bars show the distribution of click stimuli in experimental runs (considering both adaptor and target clicks) separately for each context. Symbols along the upper edge indicate the stimulus distribution mean. B. For each context, symbols represent the across-subject mean of bias in responses averaged across target locations as a function of subrun. Lines represent corresponding linear fits, i.e., temporal drifts in the responses. C. Symbols represent across-subject mean slope of the linear fit (from panel B) as a function of the stimulus distribution mean (from panel A), shown separately for each adaptor location (runs with frontal, lateral adaptor, or no adaptor). Lines show a linear fit of this relationship. Errorbars represent standard error of the mean.

The right-hand panel of Fig. A1 B shows the buildup of adaptation in response bias as a function of subrun, in a format similar to the left-hand panel. Here, the circles represent the frontal adaptor data and crosses the no-adaptor baseline data for all three response methods. The lines of different styles represent the linear fits for the different response methods, separately for the frontal-adaptor (green) and baseline (black) runs. There are clear differences between the lines for the different response methods, both in terms of their mean values and their drifts (e.g., solid lines are the most positive and decreasing, whereas the dash-dotted lines are the most negative and increasing). This confirms that a part of the drifts observed in Exps. 1 and 2 might be due to drifts in responses, not due to adaptation in the auditory spatial representations. However, important for the current study, the differences between frontal and baseline lines corresponding to the same response method always have a similar pattern, with the former having a more positive slope than the later (e.g., compare the green and black dash-dotted lines). Thus, the relative change in the slope of the drift appears to be independent of the response methods and thus may be related to adaptation in a spatial map. These results were confirmed by an ANOVA performed on the slope values with the factors of *Response method* (3 levels) and *Adaptor location* (frontal, baseline) which found significant main effects of *Response method* (F(2, 18) = 12.07, p < 0.001) and *Adaptor location* (F(1,9) = 31.3, p < 0.0005) but no significant interaction.

The right-hand panel of Fig. A1 C shows the relationship between the slope of the response drifts and the mean stimulus position for the three response methods (thin lines with different styles and the corresponding ‘o’ and ‘x’ symbols), as well as for their average (thick solid line). Consistent with the current experimental results, the average fit shows that the slope of the drift in CP is inversely proportional to the mean of the stimulus distribution (slope of this fit is −0.022; t(9) = − 7.46, p<0.0001). Importantly, the large vertical offsets between the lines corresponding to the different response methods show that the drift slopes are response-method dependent. Thus, only the relative differences obtained with the same response method (or the slopes) can be ascribed to adaptation in the spatial representation.

## Discussion

This analysis showed that the slow drift in response bias is proportional to the mean lateral position of the stimuli, independent of potential drifts in motor responses or of the environment. Specifically, stronger drifts towards midline were observed with increased laterality of the distribution mean, consistent with the idea that CP might be caused by adaptation of the neural representation to the stimulus distribution such that the neural operating points or spatial channels shift towards the stimulus distribution mean (Dahmen et al., 2010; Lingner et al., 2018). This is an alternative to a previously proposed model suggesting that repulsion-by-adaptor localization aftereffects might be caused by local suppression/fatiguing of the spatial neural channels near the adaptor caused by their extended stimulation (Carlile, 2014). While the current results are qualitatively consistent with a suppression mechanism, as the responses also drift from the adaptors, such a model does not predict that these drifts would grow with the adaptor laterality. Of course, it is possible that both suppression and shift mechanisms contribute to CP.

Note that the current analysis has several limitations. First, it assumes that the stimulus distribution mean is a relevant characterization of the distribution. Previous studies showed that other distribution statistics, like the standard deviation, also influence spatial adaptation (e.g., (Dahmen et al., 2010)). Second, it only looks at the across-target mean drift in the responses, ignoring the fact that responses for some target locations might have drifted more than others. Future studies are needed to look both other candidate statistics (e.g., stimulus variance, range, distribution median, etc.) and on the dependence of the drifts on the target location.

## FOOTNOTES

1 Note that the Appendix Figure A1 B shows the frontal and lateral adaptor run data separately.

## REFERENCES (BIBLIOGRAPHIC)

Braasch, J., & Hartung, K. (2002). Localization in the Presence of a Distracter and Reverberation in the Frontal Horizontal Plane. I. Psychoacoustical data. Acta Acust. United AC, 88, 942–955.

Brown, G. J., Beeston, A. V, & Palomaki, K. J. (2012). Perceptual compensation for the effects of reverberation on consonant identification: A comparison of human and machine performance. In 13th Annual Conference ISCA (Vol. 13, pp. 1714–1717).

Canévet, C., & Meunier, S. (1996). Effect of Adaptation on Auditory Localization and Lateralization. Acta Acust. United AC, 82, 149–157.

Carlile, S. (2014). The plastic ear and perceptual relearning in auditory spatial perception. Front. Neurosci., 8, 1–13.

Carlile, S., Hyams, S., & Delaney, S. (2001). Systematic distortions of auditory space perception following prolonged exposure to broadband noise. J. Acoust. Soc. Am., 110(1), 416–424.

Clifton, R. K., Freyman, R. L., & Meo, J. (2002). What the precedence effect tells us about room acoustics. Percept. Psychophys., 64(2), 180–188.

Dahmen, J. C., Keating, P., Nodal, F. R., Schulz, A. L., & King, A. J. (2010). Adaptation to Stimulus Statistics in the Perception and Neural Representation of Auditory Space. Neuron, 66(6), 937–948. https://doi.org/DOI10.1016/j.neuron.2010.05.018

Dingle, R. N., Hall, S. E., & Phillips, D. P. (2012). The three-channel model of sound localization mechanisms: Interaural level differences. J. Acoust. Soc. Am., 131(5), 4023–4029.

Djelani, T., & Blauert, J. (2001). Investigations into the build-up and breakdown of the precedence effect. Acta Acust. United AC, 87(2), 253–261.

Flugel, J. C. (1921). On local fatigue in the auditory system. Br. J. Psych., 11, 105–134.

Freyman, R. L., Clifton, R. K., & Litovsky, R. Y. (1991). Dynamic Processes in the Precedence Effect. J. Acoust. Soc. Am., 90(2), 874–884.

Hartmann, W. M., & Rakerd, B. (1989). On the minimum audible angle--a decision theory approach. J. Acoust. Soc. Am., 85, 2031–2041.

Herron, T. (2005). C Language Exploratory Analysis of Variance with Enhancements. January 30, 2005 version. URL: http://www.ebire.org/hcnlab/software/cleave.html.

Hládek, Ľ., Tomoriová, B., & Kopčo, N. (2017). Temporal characteristics of contextual effects in sound localization. J. Acoust. Soc. Am., 142(5), 3288–3296.

King, A. J., Parsons, C. H., Moore, D. R., & King, A. J., Parsons, C. H. & Moore, D. R. (2000). Plasticity in the neural coding of auditory space in the mammalian brain. P. NATL. A. SCI. USA, 97(22), 11821–11828. https://doi.org/10.1073/pnas.97.22.11821

Klingel, M., Kopčo, N., & Laback, B. (2021). Reweighting of Binaural Localization Cues Induced by Lateralization Training. J. Assoc. Res. Oto., 22(5), 551–566.

Kopčo, N., Andrejková, G., Best, V., & Shinn-Cunningham, B. G. (2017). Streaming and sound localization with a preceding distractor. J. Acoust. Soc. Am., 141(4), EL331–EL337.

Kopčo, N., Best, V., & Carlile, S. (2010). Speech localization in a multitalker mixture. J. Acoust. Soc. Am., 127(3), 1450–1457.

Kopčo, N., Best, V., & Shinn-Cunningham, B. G. (2007). Sound localization with a preceding distractor. J. Acoust. Soc. Am., 121, 420–432.

Kopčo, N., Marcinek, L., Tomoriová, B., & Hládek, Ľ. (2015). Contextual plasticity, top-down, and non-auditory factors in sound localization with a distractor. J. Acoust. Soc. Am., 137(EL281-287).

Kumpik, D. P., Kacelnik, O., & King, A. J. (2010). Adaptive Reweighting of Auditory Localization Cues in Response to Chronic Unilateral Earplugging in Humans. J. Neurosci., 30(14), 4883–4894.

Lingner, A., Pecka, M., Leibold, C., & Grothe, B. A. (2018). A novel concept for dynamic adjustment of auditory space. Sci. Rep., 8(1), 1–12. https://doi.org/10.1038/s41598-018-26690-0

Litovsky, R. Y., & Macmillan, N. A. (1994). Sound localization precision under conditions of the precedence effect: Effects of azimuth and standard stimuli. J. Acoust. Soc. Am., 96(2), 753–758.

Maddox, R. K., Pospisil, D. A., Stecker, G. C., & Lee, A. K. C. (2014). Directing eye gaze enhances auditory spatial cue discrimination. Curr. Biol., 24(7), 748–752. https://doi.org/10.1016/j.cub.2014.02.021

Magnusson, A. K., Park, T. J., Pecka, M., Grothe, B., & Koch, U. (2008). Retrograde GABA Signaling Adjusts Sound Localization by Balancing Excitation and Inhibition in the Brainstem. Neuron, 59, 125–137.

Moore, T. M., Picou, E. M., Hornsby, B. W. Y., Gallun, F. J., & Stecker, G. C. (2020). Binaural spatial adaptation as a mechanism for asymmetric trading of interaural time and level differences. J. Acoust. Soc. Am., 148(2), 526–541.

Phillips, D. P., & Hall, S. E. (2001). Spatial and temporal factors in auditory saltation. J. Acoust. Soc. Am., 110(3), 1539–1547.

Phillips, D. P., & Hall, S. E. (2005). Psychophysical evidence for adaptation of central auditory processors for interaural differences in time and level. Hearing Res., 202(1–2), 188–199. https://doi.org/10.1016/j.heares.2004.11.001

Recanzone, G. H. (1998). Rapidly induced auditory plasticity: The ventriloquism aftereffect. P. NATL. A. SCI. USA, 95(3), 869–875. https://doi.org/DOI10.1073/pnas.95.3.869

Shinn-Cunningham, B. G., Durlach, N. I., & Held, R. M. (1998). Adapting to supernormal auditory localization cues I: Bias and resolution. J. Acoust. Soc. Am., 103(6), 3656–3666.

Shinn-Cunningham, B. G., Kopčo, N., & Martin, T. J. (2005). Localizing nearby sound sources in a classroom: Binaural room impulse responses. J. Acoust. Soc. Am., 117(5), 3100–3115.

Stange, A., Myoga, M. H., Lingner, A., Ford, M. C., Alexandrova, O., Felmy, F., Pecka, M., Siveke, I., & Grothe, B. (2013). Adaptation in sound localization: from GABAB receptor–mediated synaptic modulation to perception. Nat. Neurosci., 16(12), 1840–1849.

Thurlow, W. R., & Jack, C. E. (1973). Some determinants of localization-adaptation effects for successive auditory stimuli. J. Acoust. Soc. Am., 53(6), 1573–1577.

Trapeau, R., & Schoenwiesner, M. (2018). The Encoding of Sound Source Elevation in the Human Auditory Cortex. Neuroscience, 13, 3252–3264.

van Wanrooij, M. M., & van Opstal, A. J. (2007). Sound Localization Under Perturbed Binaural Hearing. J. Neurophysiol., 97, 715–726.

